# Identification of biomarker and HPV strain for Cervical Cancer from pre-existing RNAseq data

**DOI:** 10.1101/355040

**Authors:** Suhan Guo, Yuhan Hao, Stuart Brown

**Affiliations:** Department of Biostatistics, College of Global Public Health; Department of Pathology, Applied Bioinformatics Laboratories; Applied Bioinformatics Laboratories Department of Cell Biology, New York University School of Medicine, New York, NY, USA

## Abstract

Human papillomavirus (HPV) increased the risk of afflicting cervical cancer. Among over a hundred strains, HPV-16 and HPV-18 caused 70% of cervical cancers and precancerous cervical lesions (WHO 2018). To reveal the profile of HPV strains, HPViewer is designed by Hao et al. (2018) for metagenomic or human genomic shotgun sequencing data analysis. The application of HPViewer in detecting HPV strains in RNA sequencing data was assessed and results were communicated in the table. The performance of HPViewer in analyzing RNA sequencing data from multiple sources, demonstrated the potential of enlarging the application of HPViewer to RNA sequencing data. Furthermore, we attempted to verify the capability of a potential biomarker p16INK4a in detecting cervical cancer from precancerous lesions. Considering the protein nature of this biomarker, the experiment was designed to detect the differentially expressed gene, associating with this protein function group, in RNA-seq data from two articles. Compare to the findings from Royse et al. (2014), confirmatory result was reproduced that comparisons between both groups yielded insignificant outcome. Since data from single article was insufficient to provide meaningful clue, final dataset was collected from multiple sources. The results were compromised by batch effect, but they supported p16INK4a to be a prospective biomarker for cervical cancer diagnosis.

## Introduction

According to Bethesda classification system, precancerous cervical lesions, also known as cervical intraepithelial neoplasia (CIN), can be categorized into Atypical squamous cells (ASC), low grade squamous intraepithelial lesion (LGSIL), encompassing CIN 1; and high grade squamous intraepithelial lesion (HGSIL), encompassing CIN 2 and 3 (Royse et al., 2014). Most precancerous lesions, particularly LGSIL, can regress due to the immune system, but some persisted and eventually developed into carcinoma. Thus, WHO guide (2013) for cervical cancer prevention advises treatment upon detection of CIN 2+, determined by cytological test - pap smear. Despite the high specificity, Pap-smear has low sensitivity in detecting the cases, and the results were not reproducible (Wentzensen & von Knebel Doeberitz, 2007). Besides, the result of Pap-smear can be obscure, rendering the follow-up tests to further determine the characteristics of atypical squamous cells of undetermined significance (ASC-US). Thus, new biomarkers were proposed to increase the sensitivity of the test. The biomarkers p16INK4a was proved to detect the underlying HGSIL with high sensitivity (Sahasrabuddhe, Luhn, & Wentzensen, 2011). Thus, after analysis of the RNA-Seq data, the oncology analysis is expected to indicate the differential expression of p16INK4a.

## Methods

Data was collected from two sources. RNA Sequencing data of three types of micro-dissected tissues, including Normal, Low-grade squamous intraepithelial lesion (LGSIL) and High-grade squamous intraepithelial lesion (HGSIL) from six patients; and thus, eighteen samples from article “Differential gene expression landscape of co-existing cervical pre-cancer lesions using RNA-Seq” was retrieved through accession ID SRP048735 (Royse et al., 2014); RNA Sequencing data of eleven patients from article “Genome-wide profiling of HPV integration in cervical cancer identifies clustered genomic hot spots and a potential microhomology-mediated integration mechanism” was retrieved through accession ID SRA189004 (Hu et al., 2015). In total, twenty-nine samples of paired-end RNA-Seq data were gathered and analyzed.

HPViewer is a tool for “genotyping and quantification of HPV from metagenomic or human genomic shotgun sequencing data” (Hao et al., 2018). This package was designed to detect HPV short DNA with less false positives by introducing two databases with different masking strategies: “repeat-mask and homology mask;” the decision to the database was made by “one homology distance matrix” (Hao et al., 2018). Following the instruction and workflow on its Github page, we prepared twenty-nine samples with “trimmomatic v0.33” to ensure homogeneity of quality (Bolger, A. M., Lohse, M., & Usadel, B. 2014). The eighteen samples from Royse et al. (2014) resulted in the average length of fifty base pairs and the eleven samples from Hu et al. (2015) had the average length of ninety base pairs. Trimmed reads were analyzed with HPViewer with average length adjusted accordingly (Hao et al., 2018).

Working protocols were recorded as Figure 1. To ensure the quality of the data, we examined all twenty-nine samples with FastQC (Andrews S., 2010) before and after the trim. Quality reports were produced. According to reports, average read lengths were stated in the previous paragraph and average quality scores of reads after trimming, were generally above twenty-eight for all samples. Paired-end fastq. files were then processed with Histat2 v2.0.5 (Kim D, Langmead B & Salzberg SL., 2015) with standard RefSeq annotation GTF file, hg19 index, to generate alignments of reads to the annotated transcribed regions of known genes; resulting in single sam. file for each. The sam.files were then compressed and sorted into bam.files with samtools v1.3.1 (Li et al., 2009). Under python environment, HTSeq-count v0.9.1 script (Anders, Pyl, & Huber, 2015) was used to count reads per gene for each sorted bam. files with reference to hg19. The output files prepared in Excel for the analysis with EdgeR package in R. After analysis with EdgeR, the first 400 gene ID, with the most significant logFC, were uploaded to DAVID Bioinformatics Resources 6.8 (Huang da, Sherman, & Lempicki, 2009) and further examined to locate target biomarker, p16INK4a. To demonstrate the differential gene expression, we generated heatmaps as presented in Figure 2.1–2.4.

**Figure 1.**
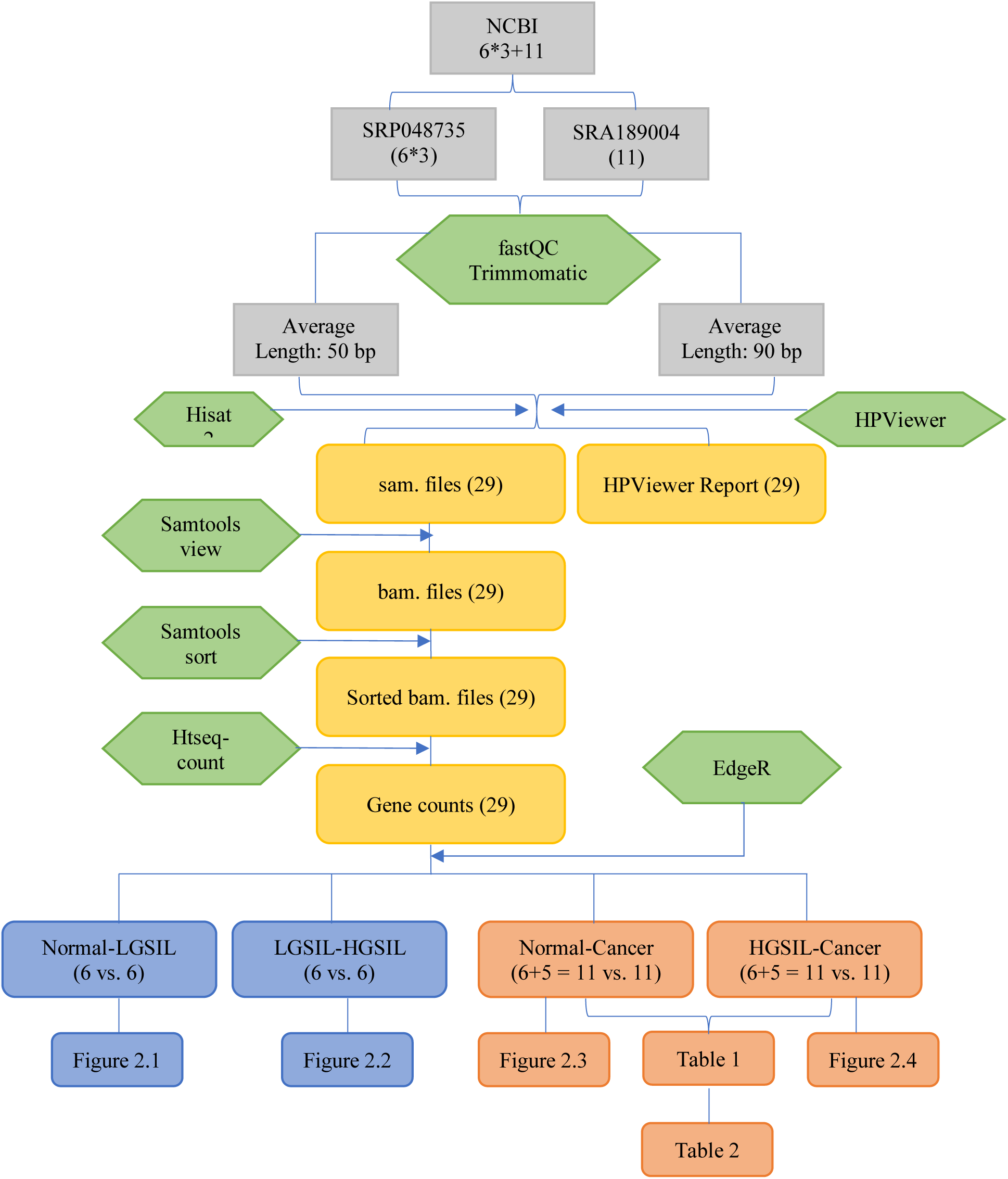

**Figure 2.1.**
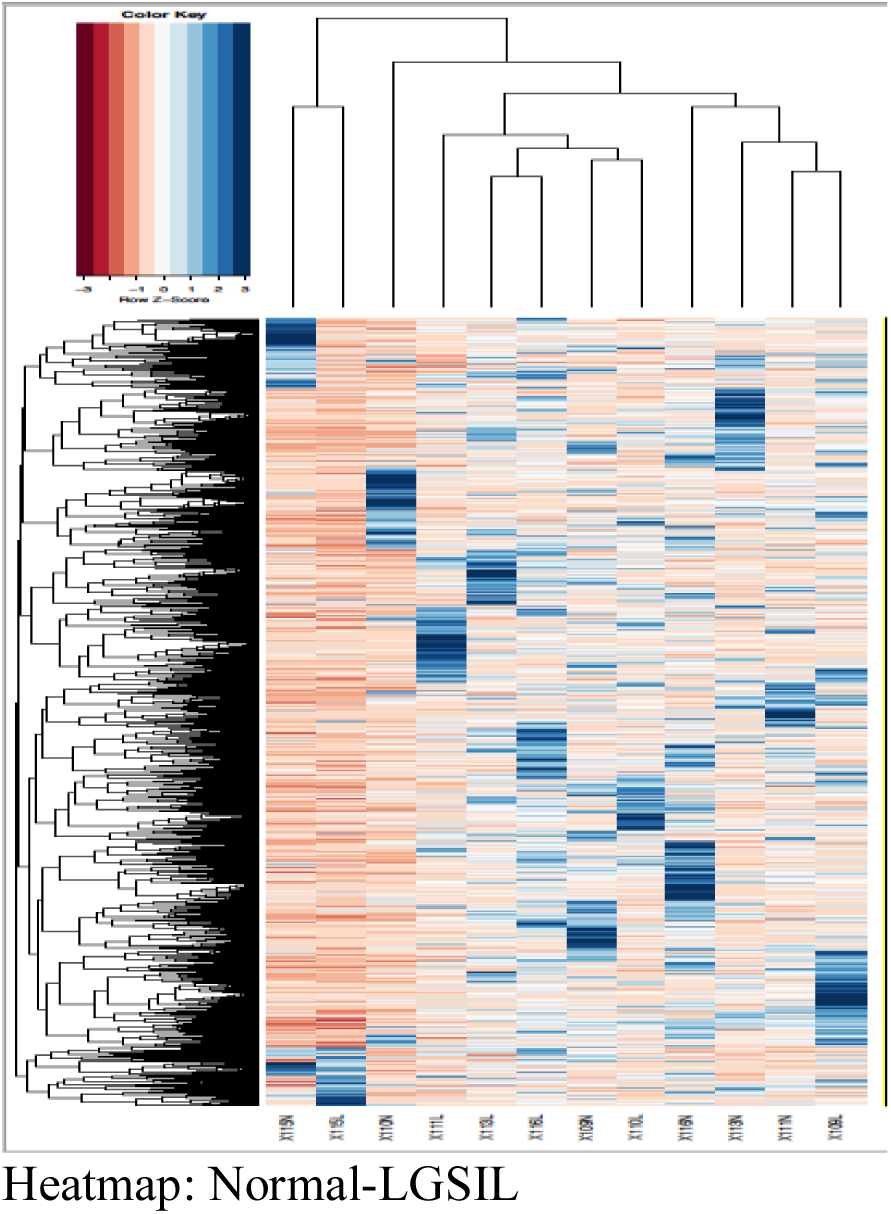

**Figure 2.2.**
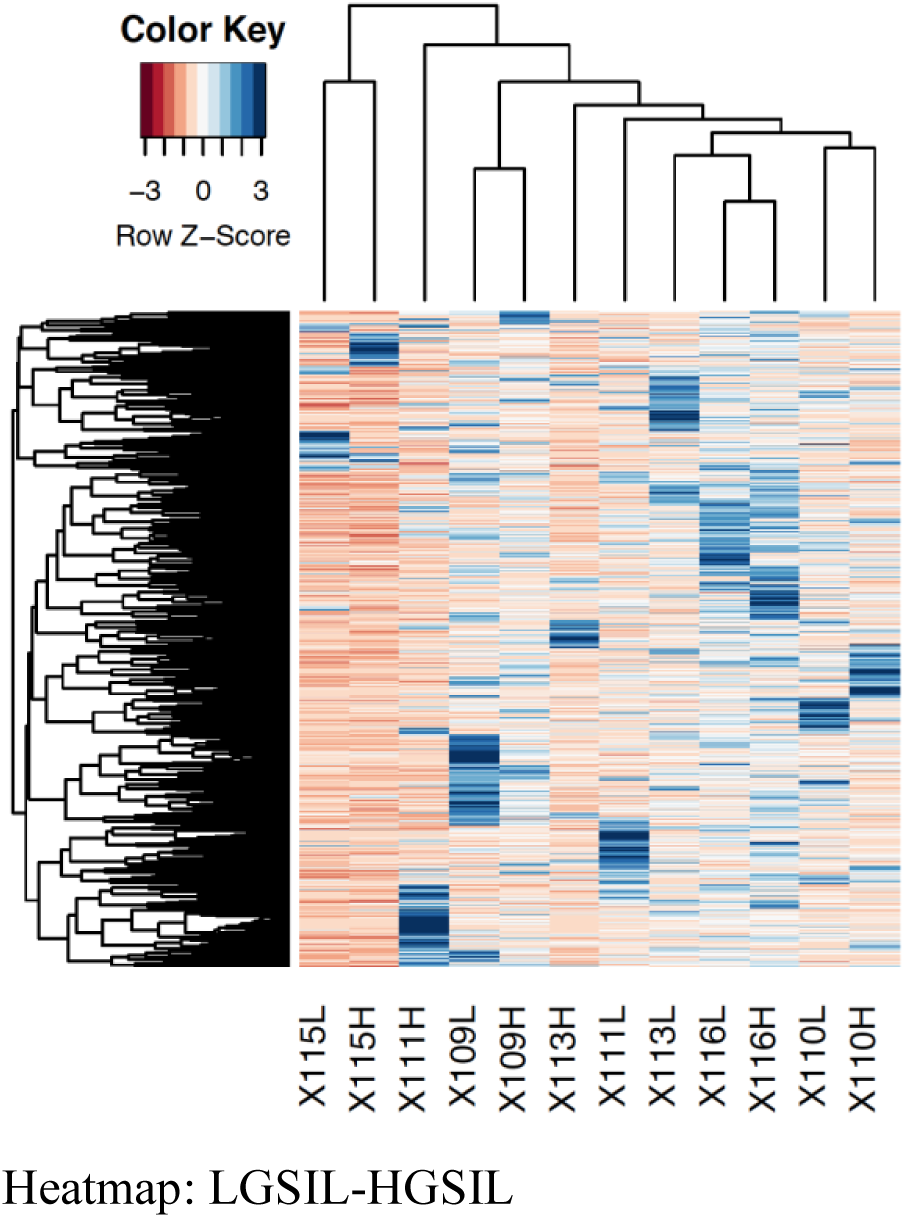

**Figure 2.3.**
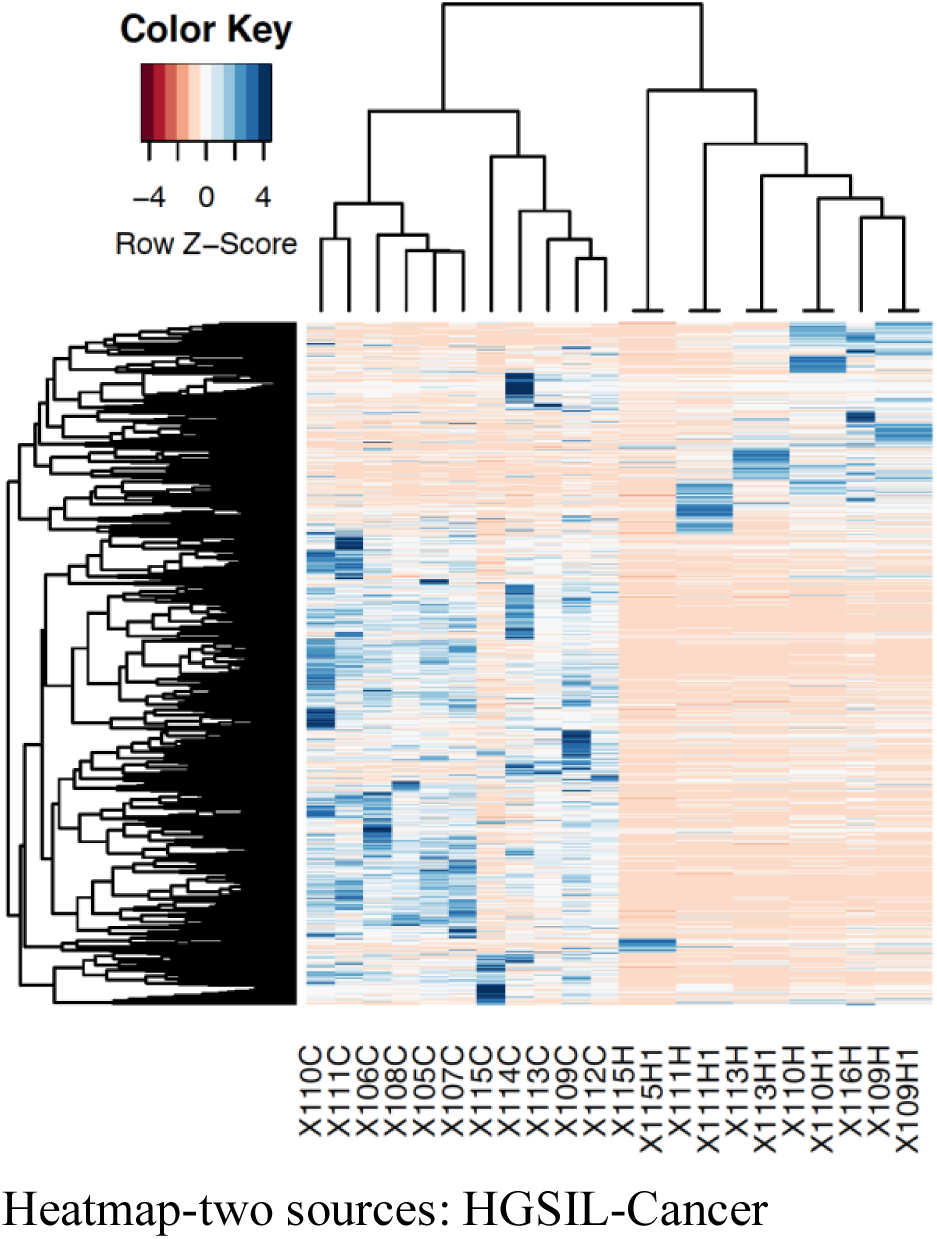

**Figure 2.4.**
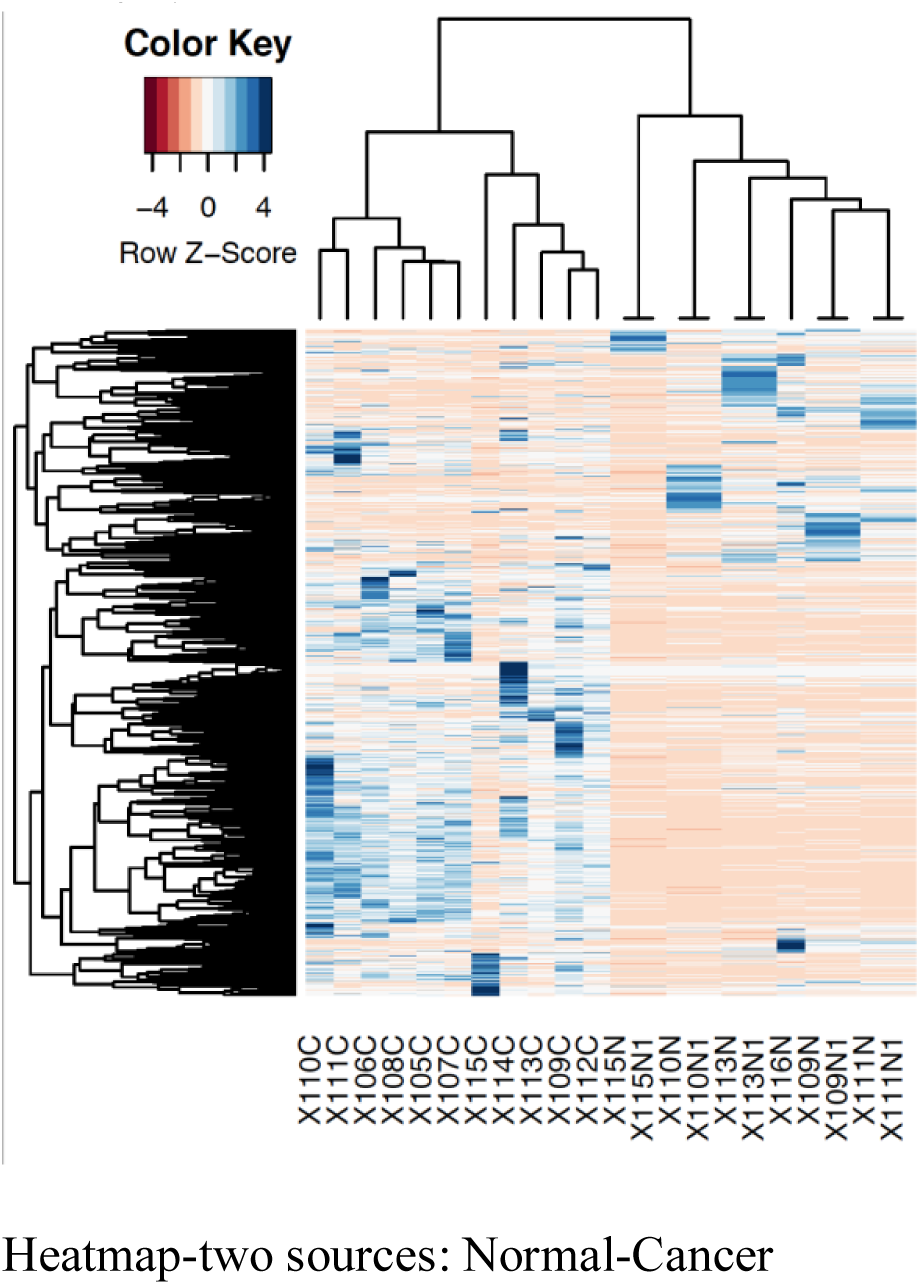

## Results

Pairing samples from all six individuals, we formulated two csv. files as instructed, which compared Normal to LGSIL as well as LGSIL to HGSIL (Royse et al., 2014). Just as mentioned in the article, no significantly differentially expressed gene was detected at the threshold: FDR ≤ .20, with these two, merged csv. files. However, comparison across articles delivered inspiring results. The eleven RNA-Seq samples of cervical cancer patients from Hu et al. (2015) were collected for validation purposes and thus, sequencing data of normal tissue, the control group, was not provided. Therefore, these eleven samples were compared to the Normal samples and HGSIL samples, with repetition respectively, from Royse et al. (2014). Diverging from our expectation, both comparisons produced significant results at threshold FDR ≤ .05: the analysis comparing eleven cancer counts to HGSIL counts detected 4479 down-regulation and 5120 up-regulation; the analysis comparing eleven cancer counts to Normal counts detected 4549 down-regulation and 5213 up-regulation as recorded in Table 1.

**Table 1.**

| **Differentially Expressed Gene counts from EdgeR analysis** |  |  |  |
| --- | --- | --- | --- |
| Comparison Group | Down-regulation | Not Significant | Up-regulation |
| HGSIL-Cancer | 4479 | 5898 | 5120 |
| Normal-Cancer | 4549 | 6039 | 5213 |

Oncological analysis of two across-article comparisons was performed in DAVID. Two up-regulated genes from Normal-Cancer comparison was found to be associated with potential biomarker p16INK4a, also known as CDKN2A in Ensembl Identification system; In HGSIL-Cancer comparison, one up-regulated gene was proved association as recorded in Table 2.

**Table 2.**

| **Differentially Expressed Gene log FC from DAVID oncology analysis** |  |  |
| --- | --- | --- |
| Comparison Group | Gene ID | Log FC |
| HGSIL-Cancer | NM_058195 | 11.45714305 |
| Normal-Cancer | NM_058195 | 7.280436839 |
|  | NM_058197 | 9.371981886 |

Among the six patients from Royse et al. (2014), HPViewer detected HPV infection in four patients and for two in four, HPV infection was detected in both HGSIL and LGSIL tissue samples. For the eleven patients from Hu et al. (2015), HPV infection was successfully detected in all the patients, and five out of eleven were infected with two strains of HPV. Results were recorded in the following Table 3.1 and Table 3.2.

**Table 3.1.**

| **Results from HPVViewer analysis on RNA-seq data: (Royse et al., 2014)** |  |  |  |
| --- | --- | --- | --- |
| **Sample ID** | **HPV infection detected** | **HPV strain** | **Count of Reads** |
| 109-HGSIL | True | HPV16 | 996 |
| 109-LGSIL |  |  |  |
| 109-Normal |  |  |  |
| 110-HGSIL | True | HPV16 | 2304 |
| 110-LGSIL | True | HPV16 | 3 |
| 110-Normal |  |  |  |
| 111-HGSIL | True | HPV31 | 1401 |
| 111-LGSIL | True | HPV31 | 4 |
| 111-Normal |  |  |  |
| 113-HGSIL |  |  |  |
| 113-LGSIL |  |  |  |
| 113-Normal |  |  |  |
| 115-HGSIL | True | HPV58 | 10 |
| 115-LGSIL |  |  |  |
| 115-Normal |  |  |  |
| 116-HGSIL |  |  |  |
| 116-LGSIL |  |  |  |
| 116-Normal |  |  |  |

**Table 3.2.**

| **Results from HPVViewer analysis on RNA-seq data: (Hu et al., 2015)** |  |  |  |
| --- | --- | --- | --- |
| **Sample ID** | **HPV infection detected** | **HPV strain** | **Count of Reads** |
| 05 | True | HPV16 | 12297 |
| 06 | True | HPV16 | 22575 |
| 07 | True | HPV16 | 26937 |
| 08 | True | HPV16 | 5008 |
| 09* | True | HPV16 | 11093 |
|  |  | HPV18 | 22 |
| 10* | True | HPV16 | 5034 |
|  |  | HPV18 | 10 |
| 11* | True | HPV16 | 4 |
|  |  | HPV18 | 43463 |
| 12 | True | HPV16 | 10189 |
| 13* | True | HPV16 | 9 |
|  |  | HPV18 | 3979 |
| 14 | True | HPV16 | 2837 |
| 15* | True | HPV16 | 1624 |
|  |  | HPV18 | 1963 |

\*More than one HPV strain was detected

## Discussion

Reproduction of results in Royse et al. (2014) was completed and confirmatory feedback was produced. With RNA-Seq analysis protocol described previously, none of the genes was verified to be differentially expressed at the pre-determined threshold. To further explore the potential of these data, we matched them to RNA Sequencing data from Hu et al. (2015). Besides the common Normal-Cancer comparison, HGSIL-Cancer comparison was designed to demonstrate the feasibility of pooling data from multiple sources; if the differential expression was proved to be absent in this HGSIL-Cancer comparison, we have the moderate confidence to deem the comparability of sequencing data from two articles. Unfortunately, this attempt failed our anticipation and thus, we have little proof to argue the variance found in Normal-Cancer comparison to be true. Instead, the significant variance can stem from several sources, such as the difference in sample collection and lab protocols. Despite all inadequacies, our analysis corroborated the potential of p16INK4a (CDKN2A) to be a biomarker for detection of cervical cancer from precancerous lesions, both low-grade and high-grade.

For a tool designed for DNA sequencing data, HPViewer performed surprisingly well with the RNA-Seq data in detecting HPV infection in precancerous lesion tissue and cancerous tissue. Supporting the description on its GitHub page, HPViewer was user-friendly due to its simplicity in command line and swiftness in producing outcome report. Since neither article revealed the HPV infection information more than a general summary, we cannot draw any conclusion on the accuracy of the tool. However, the tool proved to be sensitive to HPV strains in RNA-Seq data. Should further validation be made, the application of HPViewer on RNA-Seq data in detecting HPV strains can be deemed reliable and reproducible.

